# Detecting SARS-CoV-2 variants with SNP genotyping

**DOI:** 10.1101/2020.11.18.388140

**Authors:** Helen Harper, Amanda J. Burridge, Mark Winfield, Adam Finn, Andrew D. Davidson, David Matthews, Stephanie Hutchings, Barry Vipond, Nisha Jain, Keith J Edwards, The COVID-19 Genomics UK (COG-UK) consortium, Gary Barker

**Author notes:** https://www.cogconsortium.uk (The COVID-19 Genomics UK (COG-UK) consortium). Full list of consortium names and affiliations are available in S7 ‘*COG-UK authorship’*. Corresponding author **E-mail:** (HH).

## Abstract

Tracking genetic variations from positive SARS-CoV-2 samples yields crucial information about the number of variants circulating in an outbreak and the possible lines of transmission but sequencing every positive SARS-CoV-2 sample would be prohibitively costly for population-scale test and trace operations. Genotyping is a rapid, high-throughput and low-cost alternative for screening positive SARS-CoV-2 samples in many settings. We have designed a SNP identification pipeline to identify genetic variation using sequenced SARS-CoV-2 samples. Our pipeline identifies a minimal marker panel that can define distinct genotypes. To evaluate the system we developed a genotyping panel to detect variants-identified from SARS-CoV-2 sequences surveyed between March and May 2020- and tested this on 50 stored qRT-PCR positive SARS-CoV-2 clinical samples that had been collected across the South West of the UK in April 2020. The 50 samples split into 15 distinct genotypes and there was a 76% probability that any two randomly chosen samples from our set of 50 would have a distinct genotype. In a high throughput laboratory, qRT-PCR positive samples pooled into 384-well plates could be screened with our marker panel at a cost of < £1.50 per sample. Our results demonstrate the usefulness of a SNP genotyping panel to provide a rapid, cost-effective, and reliable way to monitor SARS-CoV-2 variants circulating in an outbreak. Our analysis pipeline is publicly available and will allow for marker panels to be updated periodically as viral genotypes arise or disappear from circulation.

## Introduction

In March 2020 the World Health Organisation characterised the global outbreak of COVID-19, caused by the severe acute respiratory syndrome coronavirus 2 (SARS-CoV-2), as a pandemic (1). A huge global effort followed to learn more about the virus, how it is transmitted and the disease it causes, in order to prevent and control outbreaks and find effective treatments and vaccines.

Since the first SARS-CoV-2 genome sequence was released in January 2020, tens of thousands of genome sequences have been shared online in public databases (2, 3). Access to sequence data is crucial for researchers to identify novel mutations, design diagnostic tests and vaccines, and to track outbreaks; allowing researchers to follow the transmission of SARS-CoV-2 both locally and globally.

As with all viruses, SARS-CoV-2 accumulates random mutations during replication. The viral replication complex has proof reading activity which may at least partially explain the relatively low rate of accumulated mutations (4). It has been estimated that SARS-CoV-2 accumulates on average about one to two mutations per month (5) which is about half the rate reported for the influenza virus that does not have a proof reading mechanism and likely has different structural constraints on its own proteins (6, 7).

Following the emergence of SARS-CoV-2, distinct lineages have formed as viruses circulating in particular regions evolved and increased in frequency. Consortia were galvanised to sequence a large number of positive SARS-CoV-2 samples to track both the evolution and geographic movements of the virus (3, 8) and a nomenclature for SARS-CoV-2 lineages was suggested to enable clear communication between research groups (9).

Contact tracing procedures that utilise genomic tools have been shown to reduce the size and duration of an outbreak (10); these tools also yield detailed information about lines of transmission. To date, SARS-CoV-2 lineages have been determined by sequencing positive SARS-CoV-2 samples. While thorough, this approach is costly and only a small proportion of positive samples have been assigned to a lineage. Our research aims to address this issue by developing a high-throughput, low-cost genotyping panel to identify circulating SARS-CoV-2 variants as genotypes (Fig 1). We use the term genotype here as opposed to lineage as our system is designed to separate samples from a local outbreak into distinct groups rather than attempt to infer their phylogenetic relationships with other samples.

**Fig 1.**
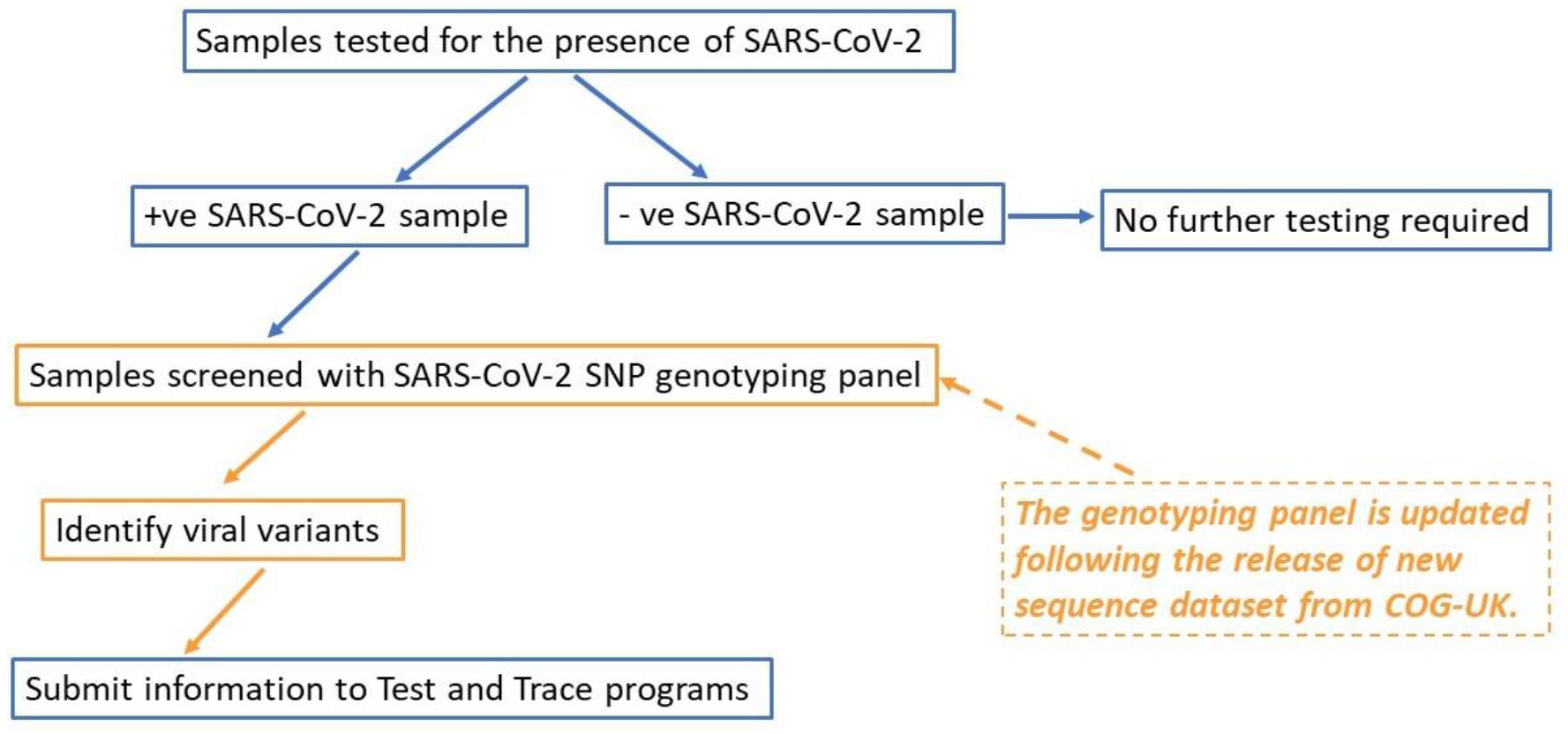
How the SARS-CoV-2 genotyping panel can be used to identify circulating SARS-CoV-2 variants.

We have validated this approach by genotyping positive clinical SARS-CoV-2 samples and show that this is an efficient method for assessing circulating variants in an outbreak.

## Materials and methods

### Samples

Extracted RNA from the supernatants of cultured cells infected with the laboratory cultured SARS-CoV-2 isolates GBR/Liverpool_strain/2020 and hCoV-19/England/02/2020 and RNA from 50 qRT-PCR positive SARS-CoV-2 samples (supplied by Public Health England (PHE) as RNA extracted from nasopharyngeal swabs) were used to validate the genotyping panel (Table 1). The hCoV-19/England/02/2020 stock contained a mixture of the wild type (wt) virus and a variant with a 24 nt deletion in the spike gene as previously described (11).

**Table 1.** Samples used to validate SARS-CoV-2 test genotyping panel. *Sample known to contain wild type and deleted spike sequences.

| Sample name | Source | Type | Sequenced | Spike Phenotype | Comparison to Wuhan-Hu-1 GenBank Acc: NC_045512.2 SNPs (amino acid substitutions) |
| --- | --- | --- | --- | --- | --- |
| GBR/liverpool_strain/2020 (GenBankAcc: MW041156.1) | University of Bristol | Viral RNA isolated from cell culture supernatant. | Yes | wt spike sequence | A6948C, G11083T, C21005T, C25452T, C28253T (nsp3: N1410T, nsp6: L37F, nsp16: A116V) |
| hCoV-19/England/02/2020 (GISAID ID: EPI_ISL_407073) | University of Bristol | Viral RNA isolated from cell culture supernatant. | Yes | Mixture* wt spike and BrisΔS | C8782T, T18488C, T23605G, T28144C, A29596G (nsp14: I150T, ORF 8: L84S, ORF 10: I13M) |
| 1 - 50 | PHE (South West Regional Laboratory) | Nasopharyngeal swabs | No | Unknown |  |

### RNA extraction

Viral RNA was extracted from cell culture supernatants using a QIAamp Viral RNA Mini Kit (Qiagen) according to the manufacturer’s instructions.

PHE samples: Viral RNA was extracted using the silica guanidinium isothiocyanate binding method (12) adapted for the ThermoFisher Kingfisher using paramagnetic silica particles (Magnesil, Promega).

### Genotyping panel design

The trimmed SARS-CoV-2 genome sequences and related metadata were downloaded from the COVID-19 Genomics UK (COG-UK) consortium website (https://www.cogconsortium.uk/data/). To check for changes in marker frequencies between May and September 2020, both the 2020-05-08 dataset (14,277 sequences) and the updated 2020-09-03 dataset (40,640 sequences) were downloaded.

### Marker selection

For SNP design, COG-UK consortium alignment data were pre-processed to select positions in the viral genome which were polymorphic with a minor allele frequency of > 0.001. After this step, sequenced accessions with identical genotypes across the polymorphic loci were removed to further simplify downstream analysis. Where two samples differed only at ambiguous base positions (no base pair called and thus recorded as ‘N’), they were considered as identical and only one was retained. Markers were then prioritised as follows. The SNP with the highest minor allele frequency was chosen as the first marker (the logic being that this allele will split the samples best into two groups). In subsequent steps, all remaining markers were evaluated to determine which one discriminated the maximum number of remaining unresolved sample pairs. The highest scoring SNP became marker 2 and the process iterated until either i) all samples could be separated into distinct genotypes, ii) no SNPs remained or iii) adding further SNPs did not result in the resolution of any additional sample pairs. For the final set of maximally informative SNPs, flanking sequences of 50 bases up and down-stream of the marker were extracted from the full sequence alignment (S1, ‘*SNPs with flanking sequence’*). If polymorphisms were observed at a frequency greater than 0.5% in the flanking sequences, they were recorded as IUPAC ambiguity codes, such that they could be avoided when designing primers for the genotyping assay. The pipeline also utilised the corresponding COG-UK metadata file to assign lineages and locations to the genotypes in our analysis output files. The complete pipeline of PERL scripts along with links to example input data files is available from https://github.com/pr0kary0te/SARSmarkers.

### Additional assays

We designed a probe set to distinguish between samples possessing the wt spike sequence and those with a known 24 nt (in-frame) deletion in the spike sequence at position 23,598 - 23,621, informally referred to as the ‘Bristol deletion’ (11), hereafter, referred to as BrisΔS (S2, ‘*Primer sequences*‘). One forward probe targets the sequence immediately prior to the deletion plus the first base of the deletion, so only gives a genotype in the absence of the deletion. The alternative forward probe targets the sequence prior to the deletion plus the first base after the deletion and only produces a genotype in the presence of the deletion. Given this design, deletions can be scored in the same way as substitutions.

### Primer design

SNP coordinates and 50 bases of flanking sequence both up and downstream of it (S1, ‘*SNPs with flanking sequences*’) were provided to 3CR Bioscience Ltd to design oligos compatible with PACE™ chemistry (13). For each of the markers in the test panel, two allele-specific forward primers and one common reverse primer were designed with a PACE-specific tail (sequences available in S2, ‘*Primer sequences*’).

### Genotyping

Genotyping was performed using the One Step PACE-RT™ (PCR Allele Competitive Extension) kit (3CR Bioscience) scaled for 1,536 plate format (the approach is described in supplementary file S3, ‘*One Step RT-PACE method*’).

Each One Step PACE-RT™ SNP genotyping reaction was performed using 2.5 ng RNA, 0.005 μL One Step RT-enzyme, 0.5 μL One Step PACE-RT genotyping master mix (3CR Bioscience) and 0.018 μL assay mix (12 μM of each forward primer, 30 μM reverse primer) in a total volume of 1 μL. The combined reverse transcription and DNA amplification reaction was performed using a Hydrocycler-16 (LGC Genomics, UK) under the following conditions: 50°C for 10 minutes; 94°C for 15 minutes; 10 cycles of 94°C for 20s, 65–57°C for 60s (dropping 0.8°C per cycle); 35-40 cycles 94°C for 20s, 57°C for 60s. Fluorescence detection was performed at room temperature using a BMG Pherastar^®^ scanner fitted with Fl 485/520, Fl 520/560 and Fl 570/610 optic modules. Genotype calling was performed using the Kraken software package version 11.5 (LGC Genomics). Fluorescent intensity was normalised for pipetting volume using the ROX standard contained within the PACE master mix.

### Data analysis

Data analysis was performed only on those samples for which 10 or more probes produced a genotype call. Samples were grouped into identical genotypes with the script qc_genotype_data.pl, which was added to the GITHUB (https://github.com/pr0kary0te/SARSmarkers) along with the SNP marker discovery pipeline.

## Results

### Minimal marker set

Up to week 18, the high-quality COG-UK sequence alignment comprised 14,277 sequences, as indicated in the accompanying metadata file. We found 41 SNPs meeting our criteria of a minimum minor allele frequency of 0.1%. Of these, our pipeline identified 22 as sufficient to provide the maximum possible discrimination between samples in the COG-UK dataset. Three SNPs were removed manually from this list as either their flanking sequences (for probe design) were overlapping or contained ambiguous bases (‘N’) close to the SNP of interest. Prior to wet-lab marker validation, we found that these 19 SNPs were capable of delineating 59 distinct variants from the COG-UK sequence alignment (S4, ‘*Regional haplotypes*’). To test the discriminatory power of the 19-marker set (hereafter, named the test set), random pairs of haplotypes for our marker positions were sampled from the COG-UK sequence alignment without replacement. We found that 89.1% of 6,202 random sample pairs were distinct at one of more marker positions. The flanking sequences for the 19 selected SNPs of the test set (S1, ‘*SNPs with flanking sequence*’), and those for the BrisΔS spike deletion, were sent to 3CR Biosciences for probe design.

### Synonymous and non-synonymous SNPs

All nineteen SNP markers in the test set target SNPs located in coding sequences. With regard to the codons within the open reading frame (ORF) of these genes, five of the SNPs were at position 1, six at position 2 and eight at position 3. Twelve of the SNPs were non-synonymous, and would result in changes to the amino acid at the given position (Table 2).

**Table 2.** Alternative SNPs and their effect on protein coding. In the Alternative Codons columns, the codon with the predominant SNP in the COG-UK 2020-05-08 dataset is listed first. Position refers to the SNP position with respect to the in-frame codon. Abbreviations: Nsp = non-structural protein; Ap = accessory protein; Non = non-synonymous, Syn = synonymous.

| Primer ID | Gene | Protein | Position | Alternative Codons |  | Syn. / Non-syn. | Alternative amino acids |  |
| --- | --- | --- | --- | --- | --- | --- | --- | --- |
| Bris_SARS-CoV-2_313 | ORF1a | Nsp2 | 3 | CTC | CTT | Syn | Leucine | ---- |
| Bris_SARS-CoV-2_1059 | ORF1a | Nsp2 | 2 | ACC | ATC | Syn | Threonine | ---- |
| Bris_SARS-CoV-2_2416 | ORF1a | Nsp3 | 3 | TAC | TAT | Syn | Threonine | ---- |
| Bris_SARS-CoV-2_2558 | ORF1a | Nsp3 | 1 | CCA | TCA | Non | Proline | Serine |
| Bris_SARS-CoV-2_2891 | ORF1a | Nsp3 | 1 | GCA | ACA | Non | Alanine | Threonine |
| Bris_SARS-CoV-2_4002 | ORF1a | Nsp3 | 2 | ACT | ATT | Non | Threonine | Isoleucine |
| Bris_SARS-CoV-2_11083 | ORF1a | Nsp5 | 3 | TTT | TTG | Non | Phenylalanine | Leucine |
| Bris_SARS-CoV-2_14408 | ORF1ab | Nsp12 | 2 | CTT | CCT | Non | Leucine | Proline |
| Bris_SARS-CoV-2_14805 | ORF1ab | Nsp12 | 3 | TAC | TAT | Syn | Tyrosine | ---- |
| Bris_SARS-CoV-2_17247 | ORF1ab | Nsp13 | 3 | CGT | CGC | Syn | Arginine | ---- |
| Bris_SARS-CoV-2_19839 | ORF1ab | Nsp15 | 3 | AAC | AAT | Syn | Asparagine | ---- |
| Bris_SARS-CoV-2_20268 | ORF1ab | Nsp15 | 3 | TTA | TTG | Syn | Leucine | ---- |
| Bris_SARS-CoV-2_20578 | ORF1ab | Nsp15 | 1 | GTG | TTG | Non | Valine | Leucine |
| Bris_SARS-CoV-2_25350 | S | Spike | 2 | CCA | CTA | Non | Proline | Leucine |
| Bris_SARS-CoV-2_25429 | ORF3a | Ap3a | 1 | GTA | TTA | Non | Valine | Leucine |
| Bris_SARS-CoV-2_25563 | ORF3a | Ap3a | 3 | CAG | CAT | Non | Glutamine | Histidine |
| Bris_SARS-CoV-2_27046 | M | Matrix | 2 | ACG | ATG | Non | Threonine | Methionine |
| Bris_SARS-CoV-2_28144 | ORF8 | Ap8 | 2 | TTA | TCA | Non | Leucine | Serine |
| Bris_SARS-CoV-2_28580 | N | Nucleoprotein | 1 | GAT | TAT | Non | Aspartate | Tyrosine |

### Evaluation of the test set

Initial evaluation of the test set and the deletion marker was performed using the two cell culture propagated SARS-CoV-2 isolates GBR/Liverpool_strain/2020 and hCoV-19/England/02/2020. The two virus genomes vary at ten nucleotide positions (Table 1) but have no differences in the wt spike gene sequences. However, in addition to the wt viral genome, the hCoV-19/England/02/2020 virus stock was known to contain a variant genome that arose during viral passage in tissue culture, which had a 24 nt in frame deletion in the spike gene sequence (BrisΔS, Table 1). Genotypes were obtained for all 20 markers (Table 3).

**Table 3.** Comparison of genotyping and sequencing data obtained for the test set and deletion marker. For the deletion marker, the ‘A SNP’ reports the wt spike sequence, the ‘T SNP’ reports the BrisΔS deletion. Sequences “Concord” where the SARS-CoV-2 isolates GBR/Liverpool_strain/2020 and hCoV-19/England/02/2020 (stock contains the wt and BrisΔS variant sequences) all share the same genotype and sequence. Separation denotes genotyping call differences between both the two isolates and the hCoV-19/England/02/2020 wt and BrisΔS variant sequences confirmed by sequencing. Alleles in the last column are those reported in the COG-UK database (from the 2020-05-08 dataset COG consortium https://www.cogconsortium.uk/data/ (14,277 sequences) with the major/minor alleles.

| Probe ID | wt Liverpool_strain |  | BetaCoV/England mix |  | Notes | COG-UK |
| --- | --- | --- | --- | --- | --- | --- |
|  | Genotype | Sequence | Genotype | Sequence |  |  |
| Bris_SARS-CoV-2_313 | C:C | C | C:C | C | Concord | C/T |
| Bris_SARS-CoV-2_1059 | C:C | C | C:C | C | Concord | C/T |
| Bris_SARS-CoV-2_2416 | C:C | C | C:C | C | Concord | C/T |
| Bris_SARS-CoV-2_2558 | C:C | C | C:C | C | Concord | C/T |
| Bris_SARS-CoV-2_2891 | G:G | G | G:G | G | Concord | G/A |
| Bris_SARS-CoV-2_4002 | C:C | C | C:C | C | Concord | C/T |
| Bris_SARS-CoV-2_11083 | T:T | T | G:G | G | Separation | G/T |
| Bris_SARS-CoV-2_14408 | C:C | C | C:C | C | Concord | T/C |
| Bris_SARS-CoV-2_14805 | C:C | C | C:C | C | Concord | C/T |
| Bris_SARS-CoV-2_17247 | T:T | T | T:T | T | Concord | T/C |
| Bris_SARS-CoV-2_19839 | T:T | T | T:T | T | Concord | T/C |
| Bris_SARS-CoV-2_20268 | A:A | A | A:A | A | Concord | A/G |
| Bris_SARS-CoV-2_20578 | G:G | G | G:G | G | Concord | G/T |
| Bris_SARS-CoV-2_25350 | C:C | C | C:C | C | Concord | C/T |
| Bris_SARS-CoV-2_25429 | G:G | G | G:G | G | Concord | G/T |
| Bris_SARS-CoV-2_25563 | G:G | G | G:G | G | Concord | G/T |
| Bris_SARS-CoV-2_27046 | C:C | C | C:C | C | Concord | C/T |
| Bris_SARS-CoV-2_28144 | T:T | T | C:C | C | Separation | T/C |
| Bris_SARS-CoV-2_28580 | G:G | G | G:G | G | Concord | G/T |
| Bris_SARSCoV2_Del_23598 (BrisΔS) | A:A (wt) | A (wt) | A:T (wt: BrisΔS) | A:T (wt: BrisΔS) | Separation | --- |

### Marker fail rate in PHE samples

The average fail rate by marker (that is, the marker produced no signal for some samples) was 18.9% ranging from 4% (marker Bris_SARS-CoV-2_25429) to 32% (markers Bris_SARS-CoV-2_2558 and Bris_SARS-CoV-2_25350). The number of fails per sample ranged from 0% (22 of the samples) to 80% (2 of the samples); those samples with less than 10 calls (8 in total) were removed from further analysis (S5 ‘*PHE 30-09-2020 genotypes*’).

### Concordance between genotyping and sequencing

The two SARS-CoV-2 isolates GBR/Liverpool_strain/2020 and hCoV-19/England/02/2020 had been sequenced, enabling a comparison with our genotyping data (Table 3). All genotyping results were concordant with the sequence data. In two cases, it was possible to confirm SNPs (at nts 11083 and 28144) differentiating the two wt SARS-CoV-2 isolates with both sequence and genotyping data. In addition, the BrisΔS sequence present in the hCoV-19/England/02/2020 stock could be discriminated from the wt sequence by the genotyping approach.

We also compared these data with the available COG-UK sequences from the 2020-05-08 dataset (representing PCR positives samples circulating March – May 2020). This showed that the majority of genotype calls concord with the major allele found in the COG-UK database.

### Genotyping clinical SARS-CoV-2 samples

To further evaluate the test set and deletion marker we genotyped 50 SARS-CoV-2 positive samples obtained from PHE (samples collected from the South West of England). For 42 of the 50 samples, results were obtained from at least 50% of the SNP markers in our panel; those that fell below this threshold were excluded from further analysis (S5, ‘*PHE 30-09-2020 genotypes.xlsx*’). For 22 of the remaining 42 samples results were obtained for all 20 markers and for a further 16 samples, results were obtained from at least 15 of the 20 markers.

We found that 12 of the 20 markers were polymorphic among the 50 PHE samples and could be used to assign them to 15 distinct groups (Fig 2 and S5, ‘*PHE 30-09-2020 genotypes.xlsx*’). To quantify the utility of our SNP panel in separating positive samples into distinct groups, we sampled random pairs of the 50 genotyped samples 1000 times and found that they were separated by at least one marker in 764 cases (76.4%).

**Fig 2.**
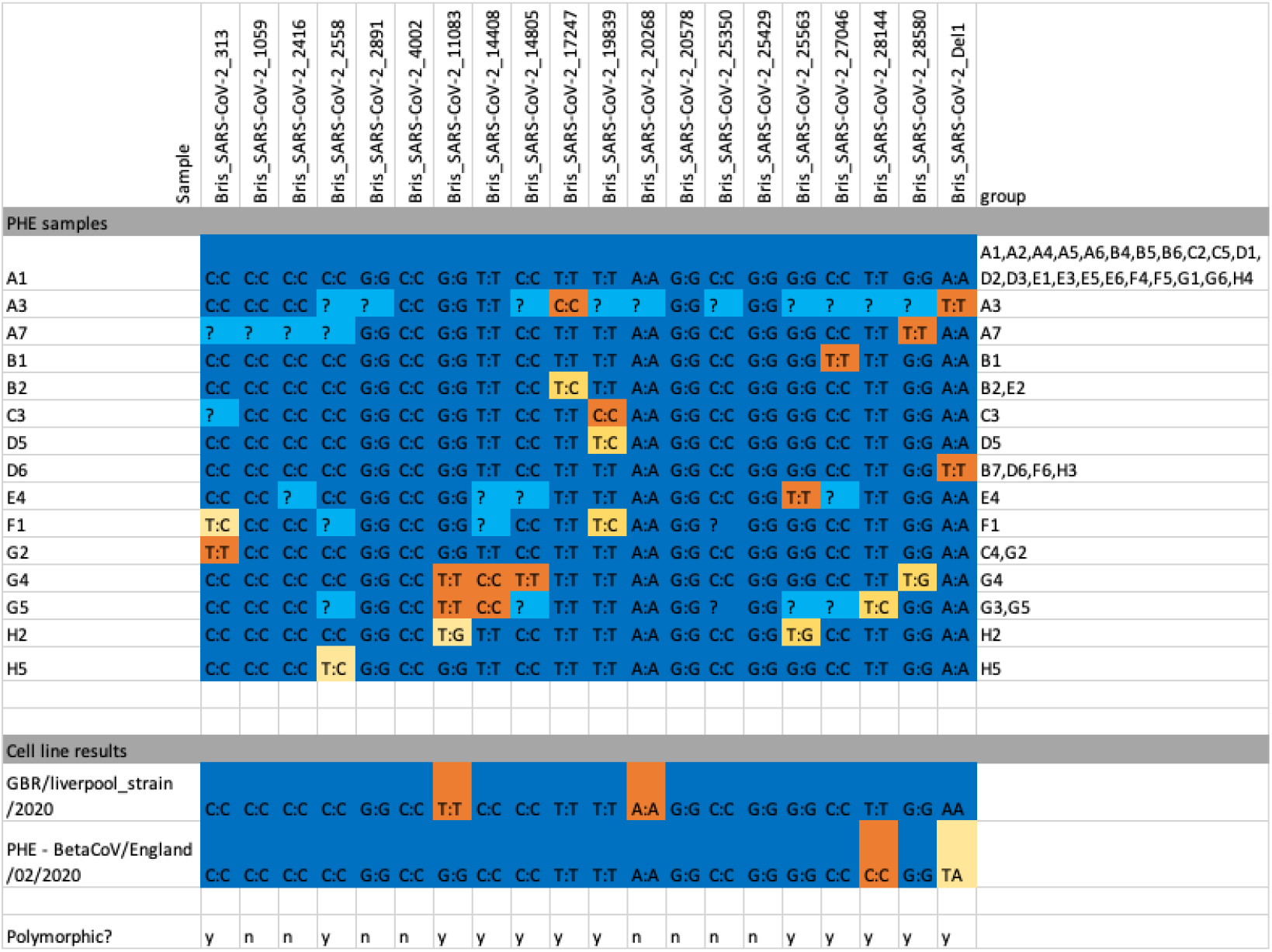
Genotyping calls for all samples. SNPs with a single allele call per sample are marked in dark blue (major allele) or orange (minor allele). Mixed calls are shown in gold and missing data in light blue. Thirteen out of 20 markers were polymorphic in our small test panel of PHE samples and cell lines and seven samples had mixed calls for one or more markers.

### Spike deletion marker

One of the markers was designed to assay a known 24 nt (in-frame) deletion, BrisΔS (11), in the spike gene (position 23,598 in the genome). This deletion has not been reported in any sequences from the COG-UK database, but we designed a probe pair in the belief that, if present, it could be detected with our genotyping panel.

The deletion marker was initially trialled with the laboratory propagated SARS-CoV-2 isolates GBR/Liverpool_strain/2020 and hCoV-19/England/02/2020 (stock contains a mixture of the wt and BrisΔS variant sequences). Illumina sequencing confirmed the wt status of the GBR/Liverpool_strain/2020 spike sequence and the mixed sequence status of the hCoV-19/England/02/2020 stock (Table 3 and Fig 2) and the genotyping data confirmed this, with RNA from the GBR/Liverpool_strain/2020 isolate producing signal only for the A base (present in the wild-type sequence) whereas RNA extracted from the hCoV-19/England/02/2020 mixed stock produced signal for both the wt A and also the T allele, which is the first base after the BrisΔS deletion (see S2, ‘*Primer sequences*‘ for details of BrisΔS deletion probes). Within the 50 PHE clinical samples assayed, seven were found to have the deletion (Fig 3a). All seven samples appeared to contain only the BrisΔS deletion and no wt spike sequence.

**Fig 3.**
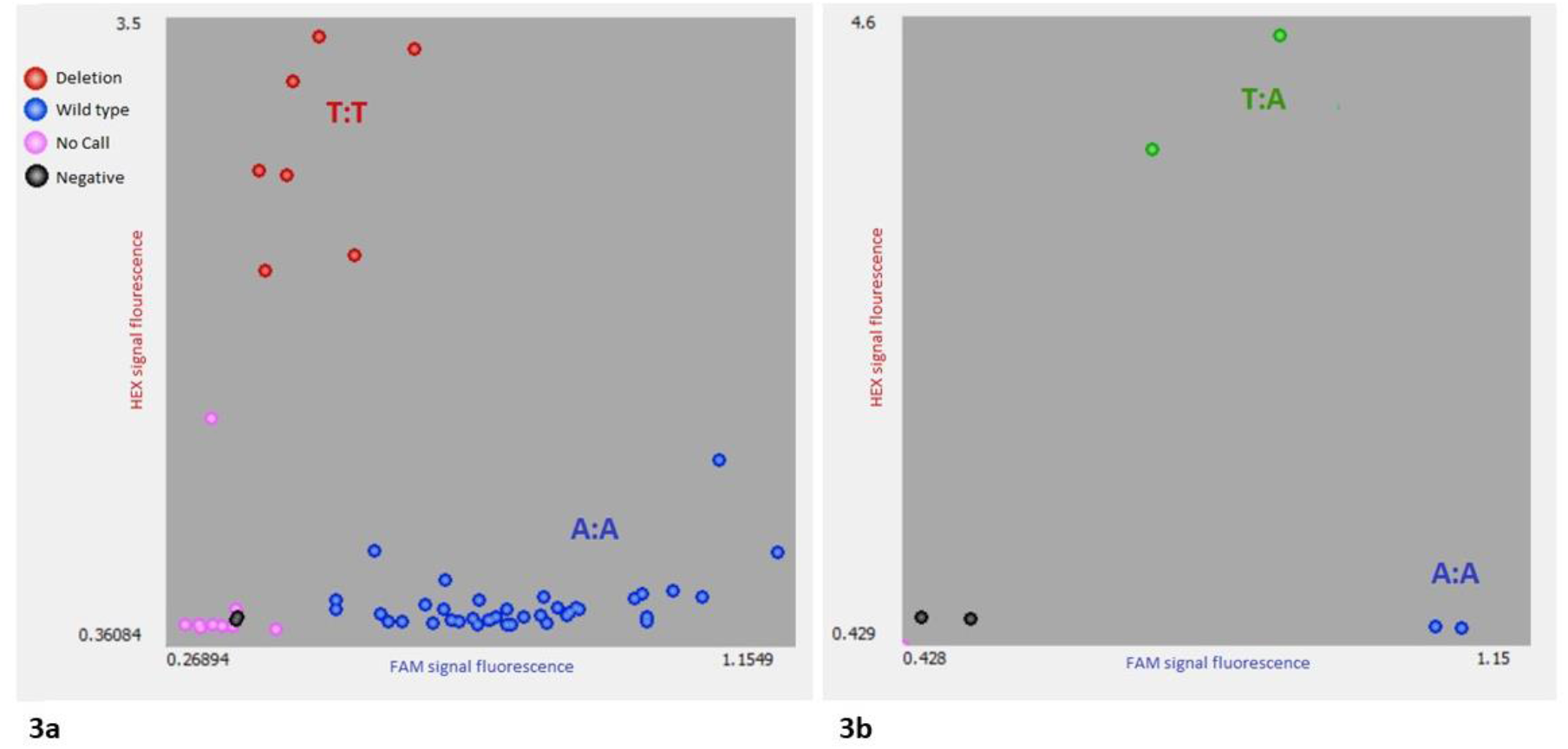
Genotyping clusters for marker BrisSARS-CoV-2_Del_23598 (BrisΔS) using PHE positive SARS-CoV-2 clinical samples (3a) and the sequenced cell cultured propagated SARS-CoV-2 isolates (3b). This marker was designed to identify the presence or absence of the BrisΔS deletion in the spike protein sequence. Sample position is determined by intensity of signal, A on the X-axis, T on the Y-axis. Unamplified samples and those between clusters were not assigned a call. Seven samples were identified with the BrisΔS deletion (shown in red).

### An evolving target

The Microreact website (14) shows how SARS-CoV-2 lineage frequencies have changed during the outbreak and similarly the SNPs we targeted in our panel also changed in frequency over time. To quantify the effect of alterations in SNP frequency over time on the discriminative power of the 19 SNP panel, it was tested bioinformatically against random pairs of samples drawn from week 19 through week 35 in the 2020-09-03 COG-UK data. The probability of the original marker set discriminating a random pair of samples decreased from 89.1 to 77.6%. There was, however, an anomaly in this analysis as our G/T SNP at position 11,083, recorded as a variant in the 2020-05-08 COG-UK data and polymorphic in our genotyping results, is reported as the non- IUPAC character “?” the 2020-09-03 COG alignment due to it exhibiting homoplasy in phylogenetic reconstruction (Andrew Rambaut, personal communication). The loss of data for this marker from the latest COG-UK alignment coupled with the absence of information on the BrisΔS deletion in the COG data means we will have underestimated the discriminatory power of our panel on more recent samples. Nonetheless, we re-ran the SNP marker discovery pipeline on the week 19-35 samples and found that the number of SNPs present at a frequency greater than 0.001 had increased from 41 to 97 (noting that the SNP at 11,083 has been masked out of that alignment) and that 51 markers were now required to discriminate all samples to the maximum amount possible. However, the majority of variants were extremely rare, such that just the first 24 markers (S6*, ‘Markers weeks 19-35’*) were capable of discriminating 95% of randomly selected sample pairs.

## Discussion

Bioinformatic analysis of COG-UK sequence alignment data from May 2020 suggested that a small number of RT-PACE genotyping assays could provide useful viral genotype identification for UK SARS-CoV-2 positive samples. We developed a genotyping ‘test panel’ of 20 markers (19 from the minimal marker pipeline plus a marker for the BrisΔS deletion). Initial evaluation of a set of two SARS-CoV-2 isolates (GBR/Liverpool_strain/2020 and hCoV-19/England/02/2020) showed that all of the markers designed produced distinct genotypes with low failure rates and comparison with available sequencing data confirmed the alleles identified in the test panel. These results were also the first demonstration of genotyping directly from an RNA virus in a single step assay.

### Clinical samples

We went on to test our panel on 50 qRT-PCR positive SARS-CoV-2 samples that were collected across the UK in April 2020. Whilst a few of the PCR-positive samples we obtained from PHE did not produce results for the majority of our marker panel, all of the markers themselves performed as expected, with missing data being attributable to low quality nasopharyngeal swabs samples rather than with any particular markers. Seven of the 20 markers were not polymorphic in the samples we were able to obtain, which was not unexpected given the small sample size. Whilst we have no reason to assume that these seven markers are not capable of producing polymorphic calls, we were unable to obtain any further samples to test this during our study. The 50 samples could be split into 15 distinct genotypes based on the genotyping data obtained and there was a 76% probability that any two randomly chosen samples from our set of 50 would have a distinct genotype. This is slightly lower than the predicted discriminatory power of the panel (89.1%) and can be explained by missing data for some sample/marker combinations, resulting from us having access to very limited quantities of PCR-positive samples, which proved to be in high demand locally for validation of qPCR assays. In a standard laboratory workflow, more RNA would be available from most qPCR positive samples.

Genotyping, unlike the reference-based sequencing, can detect mixed viral samples. We found that eight of the 50 PHE samples had mixed calls, with B2, E2, D5, G4, G5, H5 mixed at one SNP and F1 and H2 both mixed for two. We interpret this as evidence of infection by two genotypes, differing in at least one or two SNPs respectively. An example of a confirmed mixed call resulting from the presence of two genotypes was the SARS-CoV-2 laboratory strain BetaCoV/England/02/2020, which exhibited a mixed T/A genotyping call for the spike deletion and had both wt and BrisΔS deleted spike genes present in the Illumina sequence data.

### BrisΔS spike deletion marker

We hypothesised that the BrisΔS deletion at position 23,598 might be present in a subset of viral genomes in each subject and thus present as a mixed allele call. We were surprised to find that seven individuals seemed to lack the wt sequence and only possessed the BrisΔS variant. In all seven cases, the data suggest that only the deletion variant was present (unlike the mixed genotype call we observed using the hCoV-19/England/02/2020 stock). This suggests that the BrisΔS deletion variant may be capable of spreading independently of the wild-type virus. We cannot rule out the possibility that the seven deletion samples could contain a very small proportion of wt virus, but they show no evidence of this. We found no evidence of the BrisΔS deletion variant in the COG-UK alignments, which could reflect either absence of deleted samples in the database or optimisation of SNP over indel calling the COG pipeline. We also note that several deletions have previously been found in this area (15), and our primer pair will pick up any which result in the replacement of A 23,598 with T, but not others. The prevalence of the deletion and the clinical significance of this deletion therefore remain unclear and warrants further investigation. The ability of our genotyping approach to detect targeted deletions in addition to samples with mixed genotypes may prove to be useful in shedding light on the clinical significance of these phenomena.

### Panel update

A limitation of genotyping is the ascertainment bias of the probe design. Novel mutations cannot be detected which relies on an existing sequencing effort such as that performed by the COG-UK Consortium. As new mutations are discovered by traditional sequencing, the tools made available in our software pipeline may be used to design a relevant probe set for the current circulating viral population. Markers in the panel were updated based on variant analysis of the 2020-09-03 release of sequences from the COG-UK consortium to reflect the new variants circulating in the UK. We found 91 SNPs with a frequency > 0.01 in the week 19 – 35 analysis, compared to 41 SNPs in the data to week 18. The majority of the SNPs were rare, however, and we found that limiting the marker set to the most informative 24 markers gave us slightly better discriminatory power on the week 19-35 samples (95% of random pairs differentiated) than our original 19 marker set designed from week 1-18 data (89% differentiated). SNPs will continue to arise and go extinct, but our analysis suggests that a small and cost-effective panel of 20-24 markers will continue to provide useful discriminatory power in many settings.

### Application

While sequence data may offer a greater depth of information, RT-PACE genotyping can offer a rapid and low-cost solution to rapidly identify sample differences within a population. A set of 20-24 markers may be screened against 192 samples for around £2.30 per sample and savings are possible as sample numbers increase beyond this.

Genotyping is highly scalable and suited to a high throughput setting but does not require bespoke equipment which makes it suitable as an additional screening method even in smaller laboratory settings. The methods described here may be performed with only a thermocycler and FRET-capable plate reader such as that found within RT-PCR instruments. A small laboratory equipped with a 1536-well plate thermocycler and fluorescent plate-reader along with sample handling robotics and sample tracking LIMS such as KRAKEN should be able to genotype several thousand positive samples per day with input from a single trained operator.

## Conclusion

To date, SARS-CoV-2 variants have been determined by sequencing positive samples with only a small proportion of PCR samples assessed (as of 9^th^ October 2020 there were 36,593,879 reported global cases of COVID-19 and 141,000 viral genomic sequences deposited on GISAID (16). Our results show that RT-PACE genotyping with a small panel of SNPs and one indel marker can add useful genotype information to PCR-positive samples at a low cost. The fast turnaround of this approach coupled with the ease with which it can be automated means that it has the potential to provide additional detail for epidemiological studies. It is not, however a substitute for continued sequencing. Rather, the two approaches are complementary and genotyping panels will need to be cross checked against sequence alignments at regular intervals to ensure that new mutations are included and that loci which have become fixed or nearly so, are replaced. At the time of writing it is not possible to sequence every PCR positive sample in the UK and genotyping has the potential to add genotype information to all positive results with minimal investment in equipment for testing laboratories and very low cost per sample. Testing laboratories may also consider designing their own marker panels based on regional or national datasets (the latter in our case) to maximise the fit between sample SNP frequencies and the test panel. Our primer design pipeline is freely available for this purpose. The advantage of RT-PACE technology is that the SNP panel can be modified at low cost on a regular basis: in a medium to high-throughput laboratory the cost of new primer sets would not be a significant factor. The only real limitation of our approach is that it is not necessarily possible to assign samples to a specific named lineage in the way that full sequence data allows. We have shown, however that there is a high probability (>75%) of being able to separate any two samples into distinct genotypes using our marker panel, and in many settings this will be sufficient to identify or rule out transmission routes and thus inform public health policy to minimise the spread of the virus.

## Acknowledgments

This work was supported by the Elizabeth Blackwell Institute for Health Research, University of Bristol. We carried out this project in collaboration with the Bristol University COVID Emergency Research (UNCOVER) Group and we thank all of the members for their valuable feedback. We would like to acknowledge the mammoth SARS-CoV-2 sequencing effort taking place and thank the research community for making these data accessible on public databases. We are very grateful to the COG-UK sequencing consortium for making their high-quality sequence alignments and metadata available.

## Ethics statement

Samples were supplied by collaborators for the purposes of assay validation. The samples are used for the following Scheduled Purposes under the Human Tissue Act: ‘performance assessment’ and/or ‘public health monitoring’. For these purposes consent was not required under the Human Tissue Act.

## Supporting Information

**S1 SNPs with flanking sequences**

**S2 Primer sequences**

**S3 One Step RT PACE method**

**S4 Regional haplotypes**

**S5 PHE 30-09-2020 genotypes**

**S6 Markers weeks 19-35**

**S7 COG-UK authorship**

## Author Contributions

**Conceptualization:** Helen Harper, Keith Edwards, Gary Barker.

**Data curation:** Gary Barker, Amanda J. Burridge, Mark Winfield, Stephanie Hutchings, Helen Harper, The COVID-19 Genomics UK (COG-UK) consortium.

**Formal Analysis:** Gary Barker, Mark Winfield, Amanda J. Burridge.

**Funding Acquisition:** Helen Harper, Keith Edwards, Gary Barker.

**Investigation:** Helen Harper, Amanda Burridge, Gary Barker, Adam Finn, Andrew D. Davidson, David Matthews, Keith Edwards, Stephanie Hutchings.

**Software:** Gary Barker, Mark Winfield.

**Methodology:** Nisha Jain, Barry Vipond, Gary Barker, Helen Harper, Keith Edwards, Amanda J Burridge.

**Project Administration:** Helen Harper

**Resources:** Stephanie Hutchings, Adam Finn, Andrew D. Davidson, David Matthews, Nisha Jain, Barry Vipond, Gary Barker.

**Writing – Original Draft Preparation:** Helen Harper, Amanda J Burridge, Gary Barker, Mark Winfield.

**Writing – Review & Editing:** Helen Harper, Gary Barker, Amanda J. Burridge, Mark Winfield, Adam Finn, Andrew D. Davidson, David Matthews, Stephanie Hutchings, Barry Vipond, Nisha Jain, Keith Edwards, The COVID-19 Genomics UK (COG-UK) consortium.

